# STability and Shelf-Life of Plasma Bubbling Treated Cow Milk

**DOI:** 10.1101/2022.02.18.480824

**Authors:** Samarpita Dash, R. Jaganmohan

## Abstract

The demand of consumers for naturality of food with minimal processing was forced to the scientists for the discovery of non-thermal plasma which is now an emerging technology for the preservation and decontamination of highly perishable food such as milk. In this study, the microbial and physicochemical characteristics of plasma bubbling of raw cow milk were analysed and a comparison of boiled raw cow milk, commercially available pasteurised and UHT milk was observed. Again, shelf-life study was experimented for the plasma bubbling of milk with respect to raw cow milk sample (control). The plasma bubbling was generated at a voltage (200V), the flow rate of air 10 litres/hour (L/h) and applied to fresh cow milk for 5, 10, and 15 minutes (min) of time interval with 100 mL of the sample volume at room temperature. A significant reduction was observed in coliform and yeast at 200V, 10L/h, 15 min time interval of treatment. The pH of the milk was increased significantly to 6.85 after exposure to plasma bubbling. Whereas, a decreasing value was noticed in total soluble solids (TSS) and titratable acid (TA) with respect to time given by the plasma bubbling. Further a nondetrimental effect was observed for the nutrient content of plasma bubbling of milk. The result of the study indicates that plasma bubbling at (200V, 10L/h,100mL,15 min) treatment improves the quality of milk. This study shows that indirect dielectric barrier discharge (DBD) (plasma bubbling) may offer an effective microbial reduction without affecting the quality attributes. The plasma bubbling processing is an initiative for cow raw milk pasteurisation, which could have a future perspective on industrial food applications.

## 1. INTRODUCTION

Milk is a physiological liquid combination of highly biological active compounds with protein and fat^1^. Cow milk is predominant consumed all over the world in different forms and also highly perishable with rich in biological nutritive value^2-3^. Simultaneously, the hazardous effect of milk due to pathogen contamination is also well studied. Generally, the microbes present in milk are coliform, Escherichia coli, Staphylococcus aureus, yeast and mould^4^. Previous studies have analysed that the diseases caused by the contamination of milk include bovine mastitis^5^, diarrhoea^6^, milk-borne diseases found in the dairy industry^7^ due to the improper handling of dairy products. According to a previous report, per year in the USA and India, the rate of infection is 48 million due to microbes that are present in food^7^. Generally, preservation of milk evolved from chilling, boiling, pasteurization, ultra-heat treatment (UHT) process. It has been proven by studies that the pasteurization of milk can degrade the nutritional quality of milk^8^. Also, thermal processes involve many changes in the milk composition such as flavour and colloidal properties^9-10-11^. Whereas, past researchers introduced a novel and effective non-thermal technology to food science. One of the non-thermal technologies is cold plasma, used for sterilization purposes: owing to its major impact on microbes ^12-13^. The efficiency of cold plasma has been well documented in the sanitation of food^14^. The key characteristics for the deactivation of microorganisms are the UV light, source of free radicals, ions, electrons generated during cold plasma treatment. Radicals like reactive oxygen species (ROS) can directly decontaminate microbes by gaseous phase^15^. Further, the effectiveness of non-thermal plasma is applied to liquid food like milk for the inactivation of bacteria as well as for E. coli^16-17^. Recently, a study on milk sterilization was observed by dielectric barrier discharge (DBD) but the change in physicochemical properties was found^18^. Further, on low pressure plasma was observed for milk decontamination where a reduction in nutrient content such as fat was observed^19^. In contradictory to the finding, a preliminary investigation of based on plasma bubbling of milk was experimented where very negligible changes was observed for physicochemical property on milk^20^. These findings proved that the cold plasma set up playing a major role for the product quality.

Generally, cold plasma is generated by both direct and indirect methods. In direct cold plasma application: the formation of a large variety of reactive species, with a short life span (∼milliseconds) reactive species, surface plasma reactions (etching and deposition) were observed^21^. In indirect (remote plasmas), contains longer-living reactive species such as nitric oxide or ozone contact the food, while the generation of plasma was done in a separate chamber^22^. In indirect cold plasma, the quantum of heat transmission of heat to the sample is reduced. So far, an achievement for direct cold plasma has well experimented in the field of food science while the setup for indirect cold plasma is demanding^23^. Again, consumers have looked forward for dairy products which are nutritious, minimally processed, safe, healthy and economical along with a long shelf life^24^. Therefore, this study focused on a novel plasma bubbling set-up based on indirect DBD method and was established to observe the effect of bubbling of plasma on the microbial decontamination as well as the physicochemical properties during extension of shelf-life. Further, a comparative study was observed on different decontamination method such as boiled and commercialized (UHT, pasteurized) milk with raw cow milk (control) and plasma bubbling of milk for quality evaluation.

## 2. MATERIALS AND METHODS

### 2.1 Experimental set up

A cold plasma system named as plasma bubbling based on indirect DBD set up was developed^25^. Briefly, an encapsulated DBD plasma source was created using a cylindrical aluminium container named as plasma generator with two-port: one inlet and another outlet port. The inlet port was connected to an air pump through a pipe. The air pump was supplied with atmospheric gas used as feed gas and the plasma generator converted the atmospheric gas (feed gas) to free radicals. The outlet port was connected from the plasma generator to the sample through a pipe and was immersed to the sample (Fig 1). However, some modification of parameters on plasma bubbling system for generation of plasma was done in this study such as voltage, flow rate and time interval.

**Fig 1.**
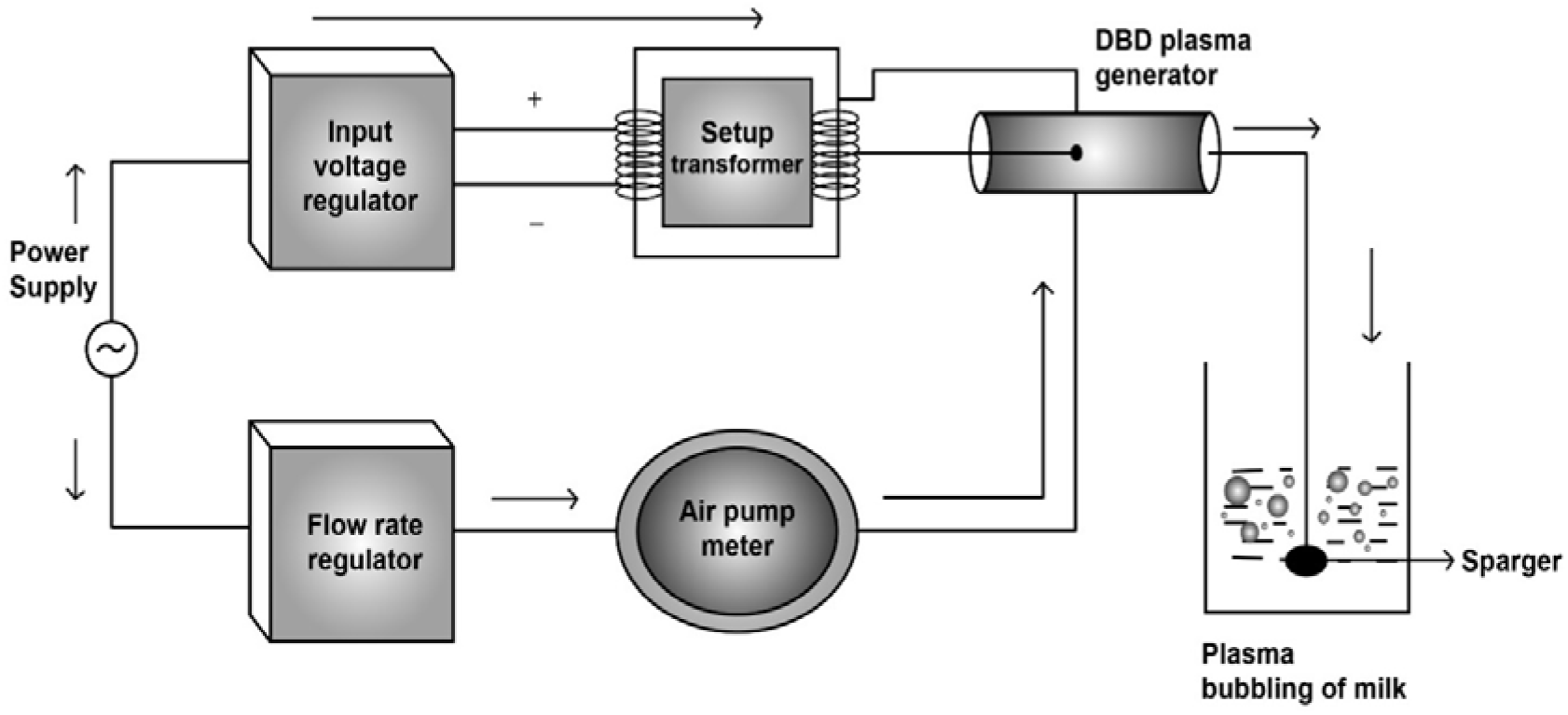
Schematic representation of plasma bubbling system.

### 2.2. Analysis of milk sample

In the present study, fresh raw cow milk collected from the local market was analysed: followed by plasma bubbling treatment at voltage 200V, flow rate of air 10 L/h, the volume of 100mL of milk sample with treatment time intervals of 5, 10 and 15 min. An extensive comparison of plasma bubbled milk was done with the fresh raw cow milk (control sample), boiled fresh cow milk at 100°C for 10 minutes^26^, commercialized (UHT and Pasteurized) milk sample.

### 2.3 Microbial analysis

The plasma bubbling of milk, control, boiled, commercialised (UHT and pasteurized) milk was tested for coliform and yeast counts by violet red bile agar^27^ and chloramphenicol yeast glucose agar^28^. Incubation was done by incubator (CI-10 plus LED Version BOD Incubator, REMI, Madurai, India) at 37ºC for coliform. While for yeast the temperature set up was 25°C^29-30^. The calculation was represented as log CFU/mL^30^.

### 2.4 pH analysis

After giving plasma bubbling treatment to the milk sample, the pH was measured and compared with the control, boiled, commercialized (UHT and pasteurised) milk using the pH meter (Laqua, Horiba Scientific, Singapore) at ambient temperature^31^ for the detection of changes in the concentration of hydrogen in the sample.

### 2.5 Titratable Acidity

Titratable acidity was performed^32^. Briefly, 20mL of the sample was taken and 5 drops of 1% w/v phenolphthalein indicator were added to the conical flask. This mixture was subjected to titration by using 0.1N NaOH and represented in % of lactic acid^33^.

### 2.6 Total soluble solids

The amount of dissolved solids present in milk was measured by total soluble solids (TSS). The TSS content was measured at room temperature (30±5ºC) by a digital refractometer (Erma, Japan, 0-80CBrix). The refractometer was calibrated with distilled water before measurement and represented as °Brix^34^.

### 2.8 Nutrient Content

The protein content is determined using protein nitrogen content of milk Kjeldahl method^35^. Fat content of milk was analysed using^36^.

### 2.9 Viscosity

The viscosity of milk was measured using Viscometer (DV-I PRIME, AMETEK, Brookfield, USA) equipped with a thermostatically controlled water. The rotor SC4-2 was used with a rotation speed of 200rpm. The measurements for control and treated samples were observed at 25±0.1°C^37^.

### 2. 10 Shelf-Life Study

The storage study of the plasma bubbled sample was compared with the control sample. All the milk samples were stored in the refrigerator at 4°C with glass jars and the samples were withdrawn once a week^38-39^ for the evaluation of important parameters of the milk i.e., analysis of microbial count, pH, TA, viscosity and TSS at the interval of 0, 7, 14, 21, 28 and 35^th^ days.

## 3. STATISTICAL ANALYSIS

The result was analysed by using SPSS version 22 statistical software (SPSS, Inc., United States). All the experimental analyses were replicated three times. The statistical analysis was carried out by using a one-way analysis of variance (ANOVA). When significant deviations were detected, the differences among the mean values were determined by executing Duncan’s multiple tests of comparison at a confidence level of p< 0.05.

## 4. RESULT AND DISCUSSION

### 4.1 Microbial Study

The result of viable cell count of coliform and yeast was monitored in the plasma bubbling, control and boiled milk sample which was compared with commercially available UHT and pasteurized milk samples (Fig 2). Plasma bubbling treated milk sample for 5, 10 and 15 min at (200V, 10L/h,100mL) was mentioned as M1, M2 and M3 respectively. A statistically reduction (P<0.05) i.e., no viable cells were found for M3 treated sample while the control value was 6.38±0.002 and 4.42±0.002. These results indicate that the plasma bubbling system used in this study can reduce the number of microbes in milk with an increase of interval of time. A significant reduction was observed in coliform, yeast after plasma bubbling. While the treated sample M3 was able to decontaminate microbes from cow raw milk samples successfully. This could be explained due to the development of reactive species during plasma generation: thus, showing a better efficacy on microbial cells. Previous studies reported that with an increase in time and flow rate in plasma bubbling, the hydroxyl radical (^•^OH) generation was increased^26^, which could be responsible for the destruction of microbial cell^40^. Again, the commercialized UHT and pasteurized milk decontaminates from all microorganisms, but already it has been studied that the physical properties were affected during the shelf-life study of UTH and pasteurized milk^9-8^.

**Fig 2.**
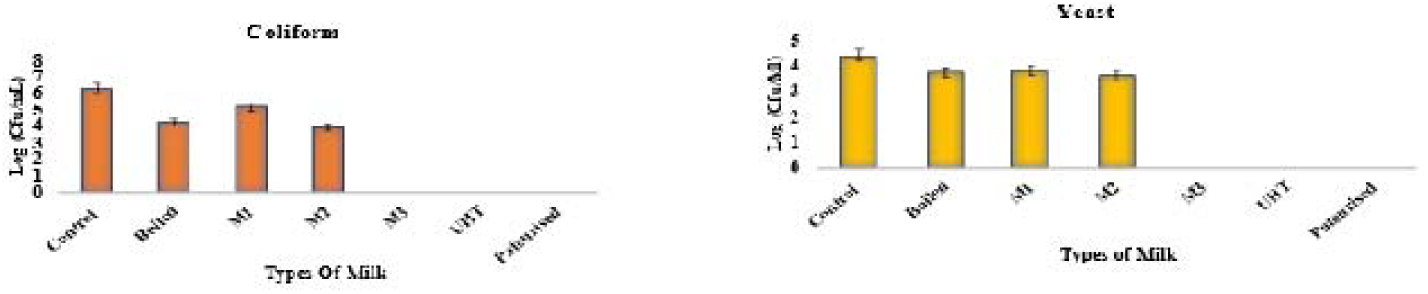
A comparative study on microbial cell viability of plasma bubbling treated milk with control, boiled, Commercialized (UHT and pasteurised milk).

The time interval of 15min was taken as the final treatment time because of its satisfactory deactivation of microbial load. Specifically, in dairy products, the result of cold plasma treatment on microbes depends on various factors such as target species of microorganism, time interval, input power, gas and food composition^41^. The inactivation of microbes by using cold plasma was not fully observed since the nature of plasma is highly complex ^42^.

Basically, the acceptable range of coliform in milk is 0-1000/ mL at 24h with 37°C^43^ which was owned by the plasma bubbling treated sample M3. Previously it has been observed that the yeast cells are found in both raw and pasteurised milk ^44-45-46-47^ but at a low population mostly 10^3^ cells /mL^48^, while in this study at M3 treated sample yeast cells were not observed which indicates a good finding.

### 4.2 pH

The hydrogen concentration is the most important factor in milk. The control value of pH was 6.66 ± 0.015 (Fig 3). While a statistically significant (P<0.05) increase was observed in pH after the application of plasma bubbling. The pH value 6.85±0.01 in M3 which is showing a gradual increase with respect to the time. This could be due to the increase in hydroxyl **°**OH radicals’ generation during cold plasma generation^49^. A contradictory result was found in cold plasma treated milk^19^. The result obtained for boiled milk samples showed decrease value in hydrogen concentration which could be explained by the changes in gelation behaviour during heating of milk^50^.

**Fig 3.**
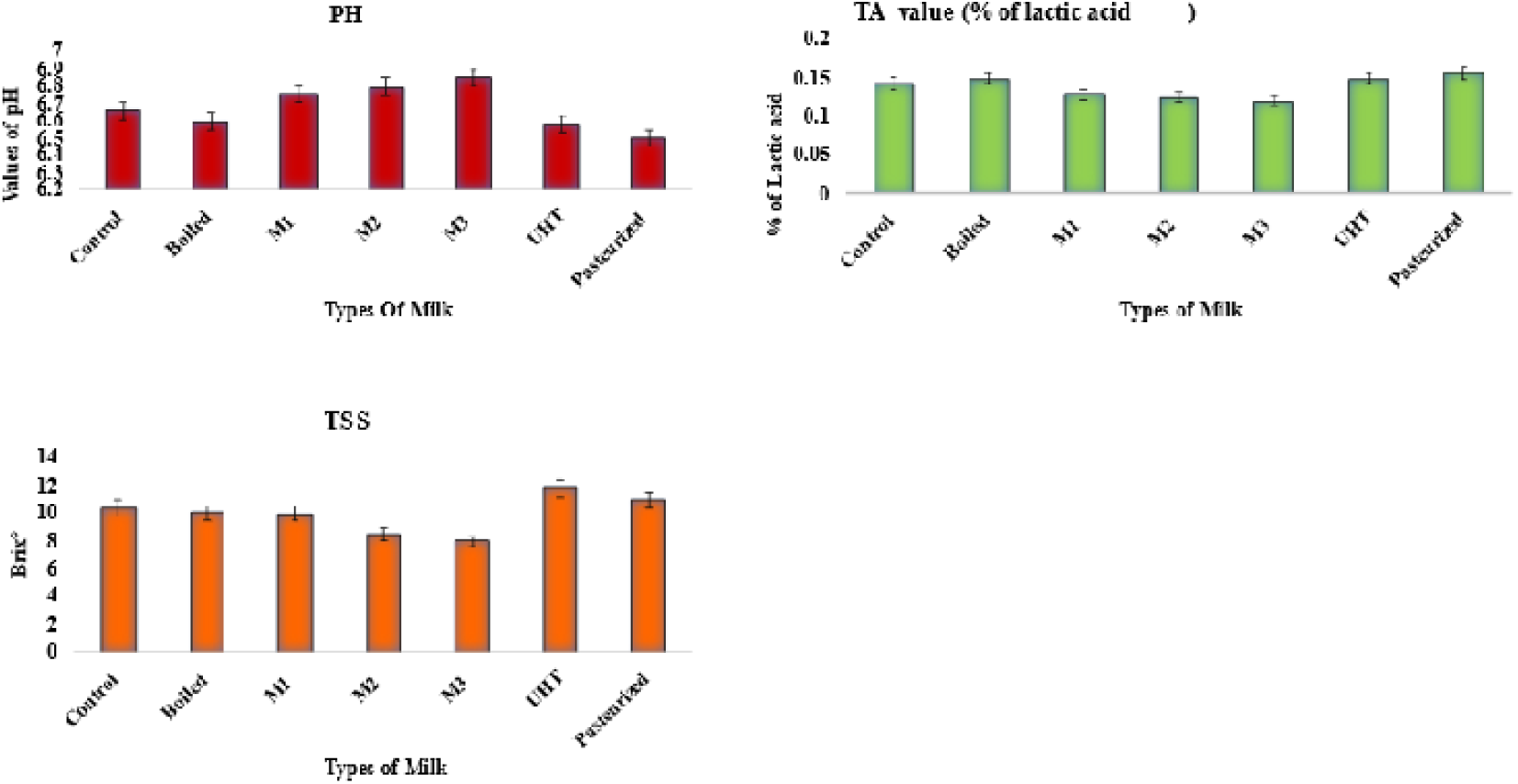
A comparative study on physicochemical property of plasma bubbling treated milk with control, boiled, Commercialized (UHT and pasteurised milk).

### 4.3 Titratable acidity

In the post plasma bubbled milk sample with respect to the time interval, a significant decreasing value of % of lactic acid was observed at M3 treated sample i.e., 0.121±0.002 (Fig 3), whereas the control sample TA value was 0.144±0.002. This could be explained due to the increase of OH^•^ radical formation during cold plasma treatment which is responsible for the decomposition of an additional water molecule into the sample^51^. While, a contradictory to this study an increase in TA value was observed in tiger nut milk after exposure to cold plasma^32^. The boiled milk showed significant (P<0.05) increase in TA value 0.15±0.002. The increase in TA value of boiled sample could be due to the high heat treatment which was responsible for the degradation of lactose^52-53^.

### 4.4 Total soluble solids

The value of TSS was decreased after a long time of exposure to plasma bubbling. The value of °Brix of control sample was 10.4±0.003 while at M3 treated sample the value decreased to 8±0.001. This decrease in °Brix value could be explained due to the ozonolysis, happening during plasma exposure which made cleavage of the glycosidic bond and helps to de-polymerisation of macromolecule^54-55^. A comparative description was mentioned for the physicochemical property of milk (Fig 3). This contrast, previous study revealed that: no significant change was observed in tiger nut milk after expouser to cold plasma^32^. The TSS value of the boiled sample is non significantly decrease (P <0.05) 10.1±0.001. The slight decrese in TSS could be due to evaporation of water from milk during boiling.

### 4.5 Nutrient content

The protein and fat content of the raw milk (control), plasma bubbling treated sample, boiled sample along with commercialized (UHT and pasteurized) sample is given in the (Fig 4). Protein and fat content in milk after plasma bubbling treatment for all the treatment time i.e., (M1, M2 and M3) were found to be significantly not different (P>0.05) from the control. For protein and fat content the value of control sample was 3.41±0.001% and 3.52±0.001% respectively. While, for M3 the value of protein and fat content was 3.39±0.002 and 3.49±0.001% respectively. A similar findings of protein content after cold plasma was observed on milk^56^. Further, in a recent study, a nondetrimental effect was observed on cold plasma of milk^19^. In the present study the fat content of milk was slightly decrease which could be due to the colour difference on milk after exposure to cold plasma due to the reactive species which have high oxidation ability^57^. A similar finding on fatty acid composition of milk was observed^58^. A non-significant (P<0.05) increase value was observed for the boiled sample for fat content 3.53±0.003. The negligible increase in fat content of boiled sample could be due to the loss of evaporated water during heat treatment^59^. While a decrease in protein content was observed after boiling of milk could be due to heat which was responsible for the denaturation of whey protein particularly beta lactoglobulins^60^.

**Fig 4.**
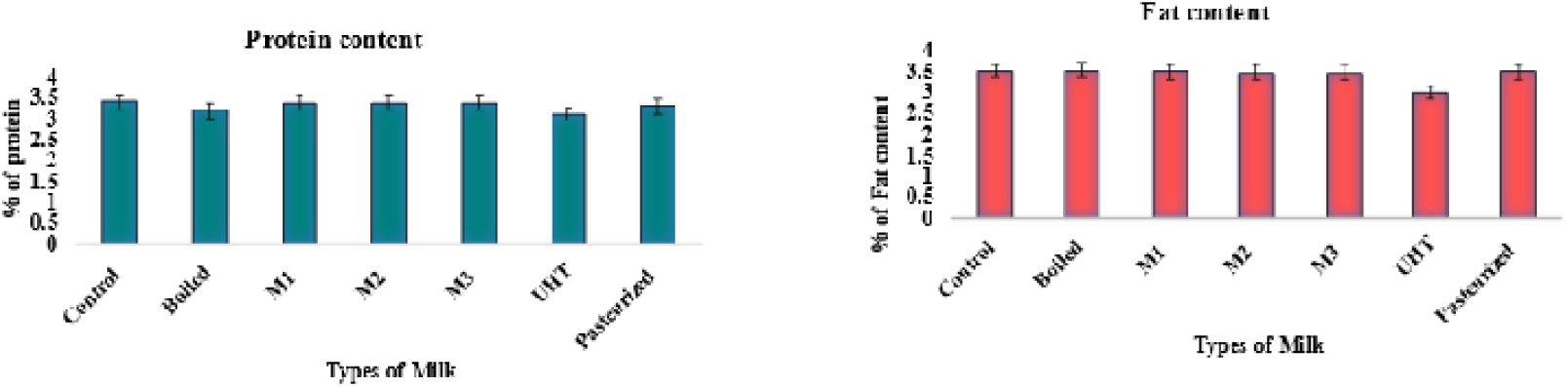
A comparative study on nutrient content of plasma bubbling treated milk with control, boiled, Commercialized (UHT and pasteurised milk).

### 4.6 Storage study of milk

In the present study, a comparison of storage study was done for plasma bubbled milk i.e., 200V,10 L/h, 100mL at time interval of 5, 10 and 15 min which compared with the control sample (Table 1). Plasma bubbling is a non-thermal technology based on indirect DBD, where the treatment used to carry out at room temperature due to this reason control sample was compared with the plasma bubbled milk sample. The shelf life of the plasma bubbled milk samples was established by taking into account the initial microbial populations of the control sample. During storage, there were an increase of coliform and yeast counts in the control sample was observed and spoilage was declared on the 3rd day of storage. The coliform and yeast count of the control sample were 6.38 and 4.42 respectively, for the 0^th^ day. During storage period in the plasma bubbling treated sample: a drastically reduction was observed on microbial load for M1, M2 and M3 (Fig 5). The sudden reduction of microbial cell could be described by the hydrogen peroxide produced by cold plasma^61^ which is stable and storable at a low temperature ranging from (−60-0ºC)^62^ Further, hydrogen peroxide is considered as an antimicrobial compound^12-63^. A similar finding of sudden decrease in microbial load was observed on cold plasma of milk^16^.

**TABLE 1.**
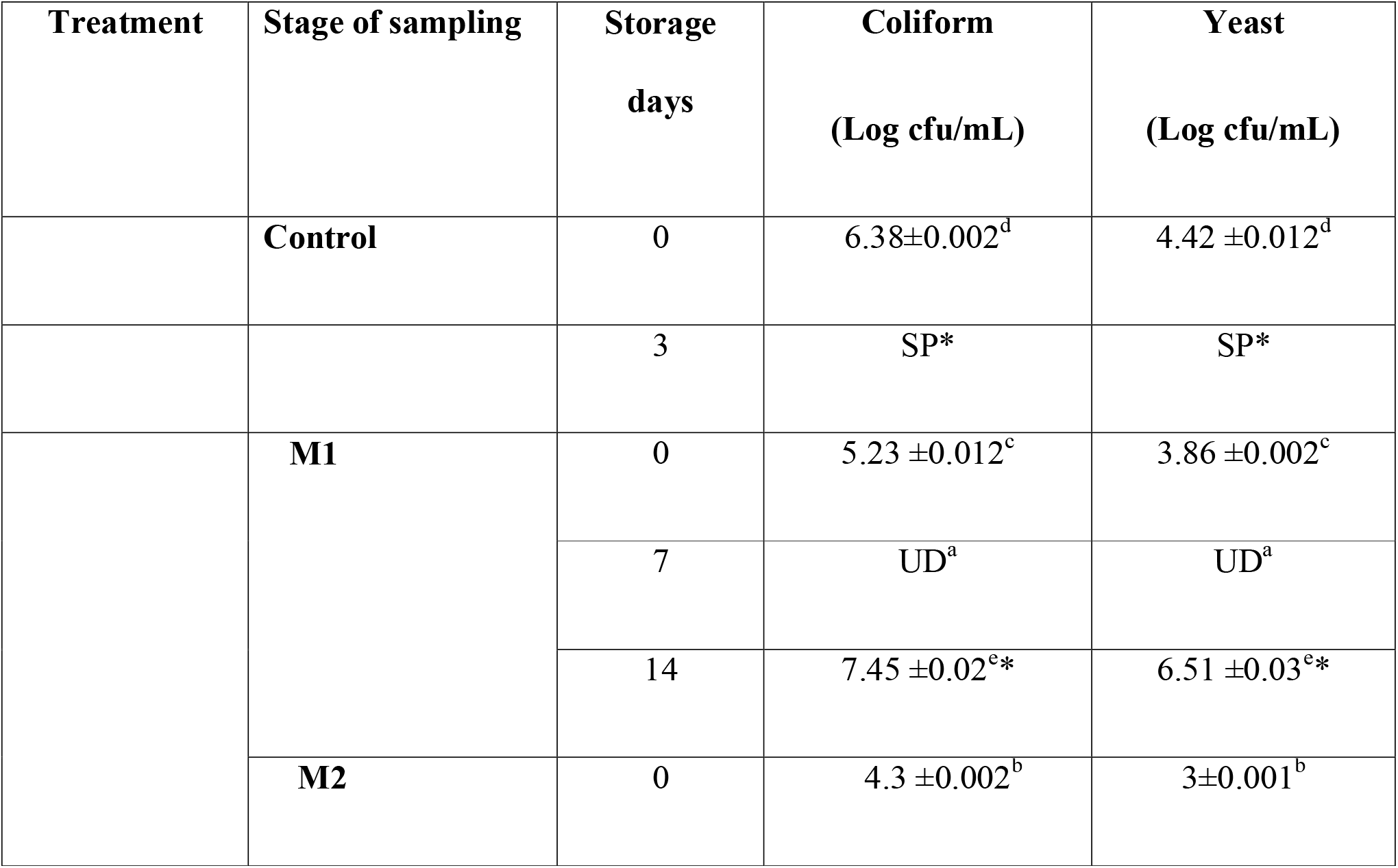
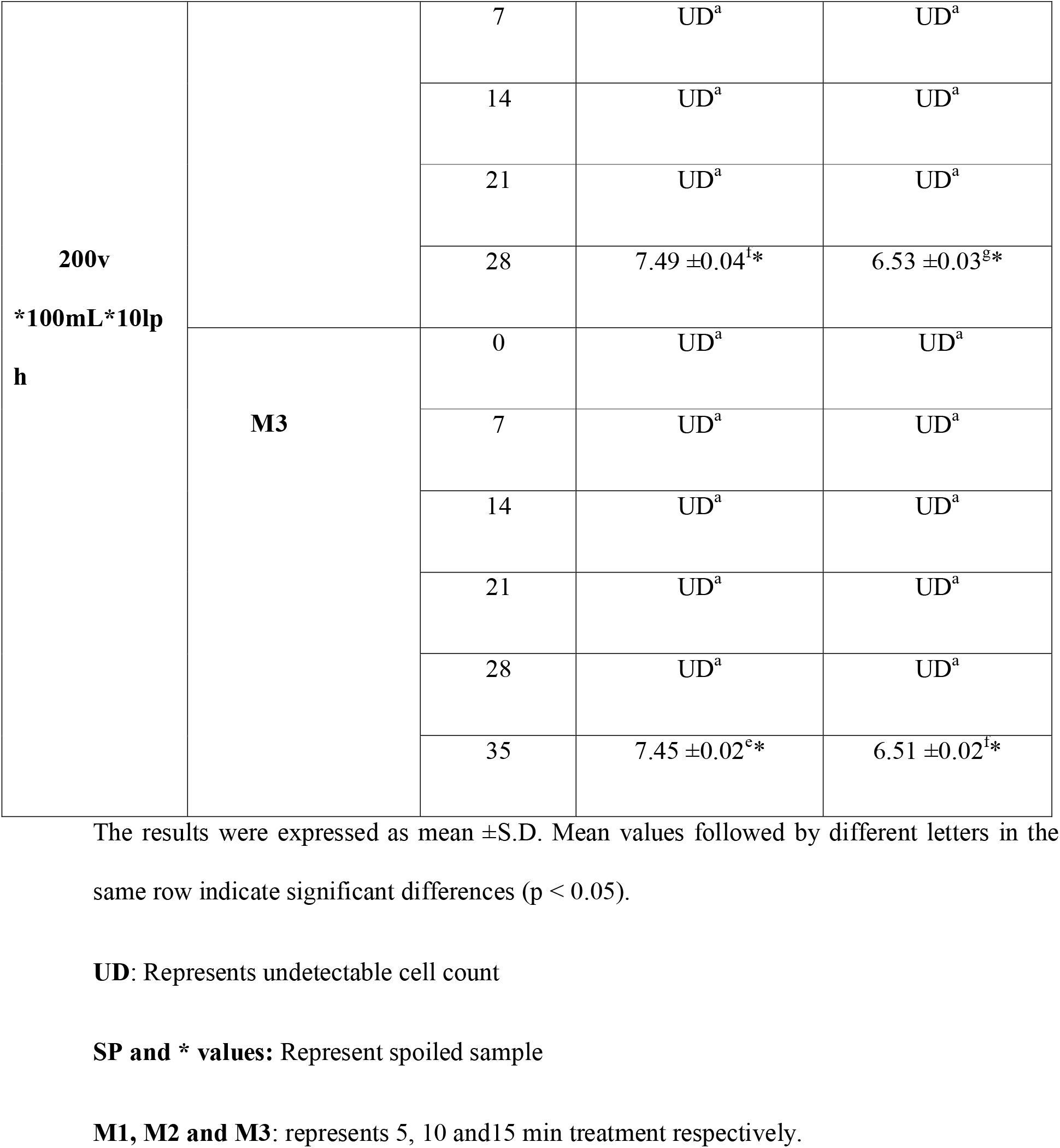
Microbial Analysis during storage

**Fig 5.**
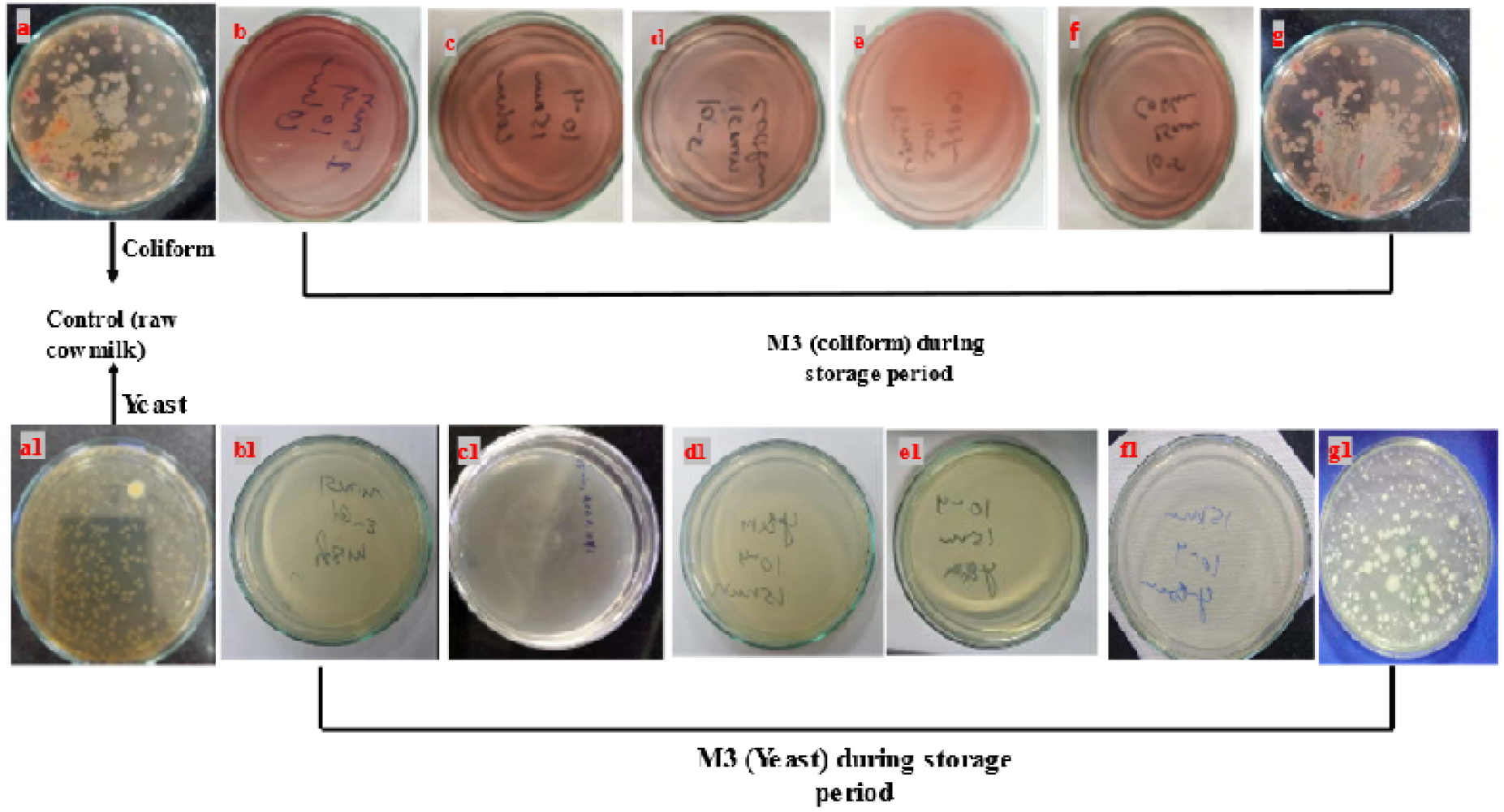
A comparative study on coliform viable cell count in raw milk and M3 (15 min, 10L/h, 200V) treated sample during storage period a) control (untreated), b) M3 on 0^th^ day, c) M3 on 7^th^ day, d) M3 on 14^th^ day, e) M3 on 21^st^ day, f) M3 on 28^th^ day and g) M3 on 35^th^ day (spoiled). For yeast viable cell count in raw milk and M3 treated sample during storage study a1) control (untreated), b1) M3 on 0^th^ day, c1) M3 on 7^th^ day, d1) M3 on 14^th^ day, e1) M3 on 21^st^ day, f1) M3 on 28^th^ day and g1) M3 on 35^th^ day (spoiled).

The spoilage was calculated based on the activation of microorganisms. The milk sample treated for M1 was spoiled on 14 days of storage while for M2 and M3 the spoiled was declared on the 28^th^ day 35^th^ day of the exposure respectively, the spoilage of milk could be described by activation of coliform which helps in the fermentation of lactose by producing of gas and acid^64-63^.

### 4.7 Physiochemical properties of stored milk

During storage, for the control milk: the pH decreased to 4.2±0.013 on 3^rd^ day of storage; whereas a gradual increase in pH was observed for the treated sample on the 7^th^ day of storage for M1, M2 and M3 plasma bubbling treated sample. The sudden increase in pH could be due to the OH**°** radical production in cold plasma technology^49^: the closed glass vessel could maintain the OH^•^ radical formed on the cold plasma generation during storage at refrigerator condition. However, the M1 spoiled on the 14^th^ day of storage, M2 on the 28^th^ day and M3 on 35 days of storage with pH value 4.1±0.015, 4.0± 0.012, 4.1±0.015 respectively, the decrease could be explained due to the acidification of milk^65^ (Table 2). The spoilage of milk was calculated based on the activation of microorganisms and pH value. The other parameters of physicochemical properties were also observed during the storage period in which % lactic acid increased with respect to the treatment time. The value of % lactic acid for M3 was 0.144±0.002 on the 35^th^ day of storage while control showed a value of 0.202±0.013 on the 3^rd^ day of storage: the increase in % lactic acid could be explained due to the high bacterial activity during storage^65^. Again, TSS (°Brix) increased significantly with progress in storage days, which could be owing to the hydrolysis of components ^66^. Further, the viscosity was observed for M1, M2 and M3 with respect to control sample. In the initial day (0^th^ day) the viscosity of the plasma bubbling treated milk statistically showed a non-significant value i.e., negligible decrease. M1, M2 and M3 value was 1.66, 1.65 and 1.64cP respectively, while the value of control was 1.67cP. The negligible decrease in viscosity value could be postulated due to high voltage during cold plasma generation which is responsible for the oxidation of protein and lipid^67^. A contradictory finding was observed on cold plasma of milk^68^. Again, during storage days the value of viscosity remain constant which could be due to lower temperature specifically at 2-5°C on refrigerator condition which was able to deferred the time for viscosity change and responsible to maintain water retention ability of macromolecular substances in milk. Eventually, an increased value was observed during spoilage of milk could be due to the denaturation of casein^37^. However, the control milk shelf life was of 3 days while for plasma bubbling treated milk the shelf life was 14, 28 and 35 for M1, M2 and M3 respectively.

**TABLE 2.**
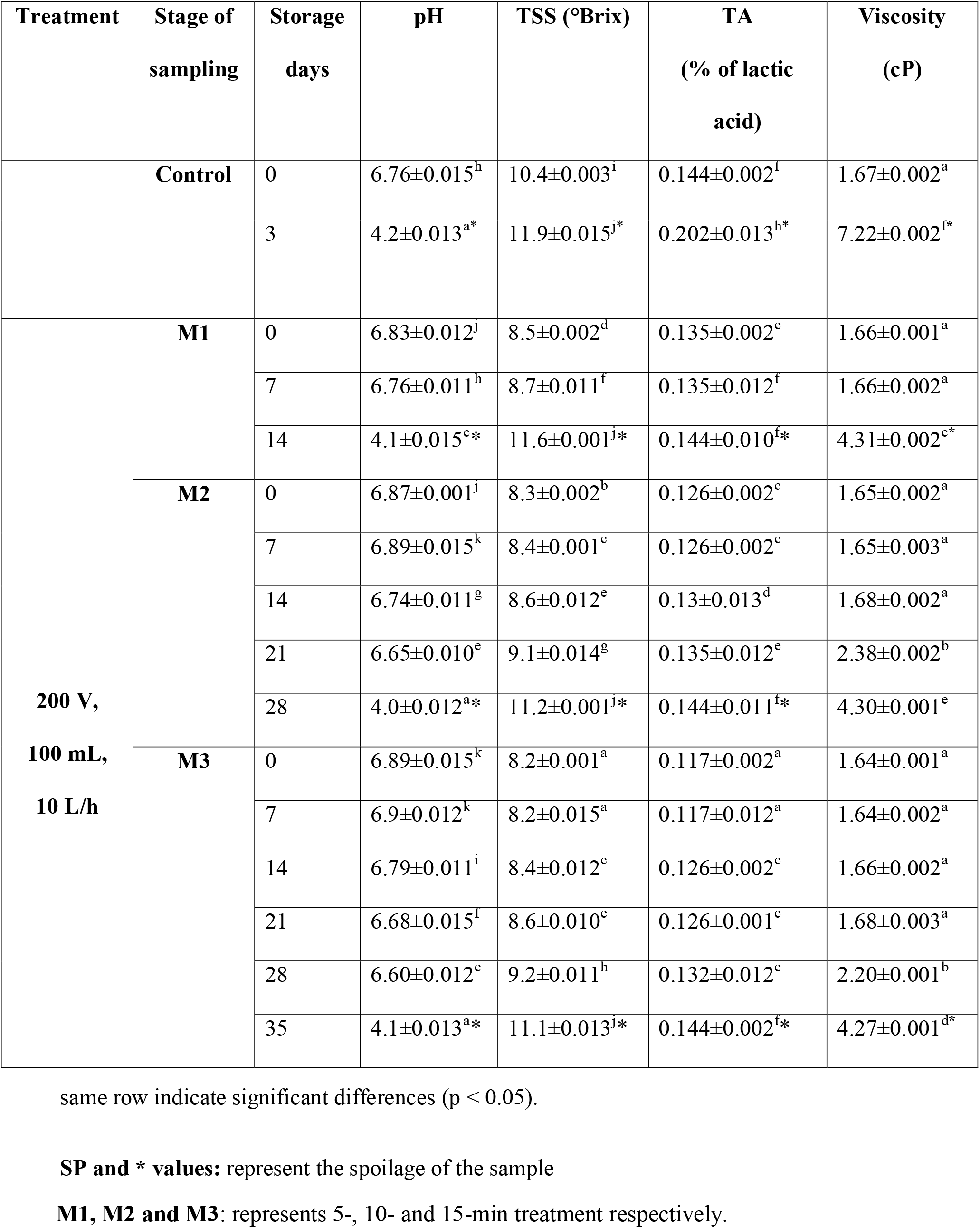
Physiochemical property of storage study

## 5. CONCLUSION

Next to natural raw cow milk the quality of the milk is maintained by cold plasma technology introduced to food industry. The freshness, safety and natural characteristic are observed in plasma bubbling. So far, no other technology has been produced favourable property along with microbial safety concerns simultaneously. The plasma bubbling system was able to decontaminate microbes like coliform, yeast satisfactorily. The treatment time was played a major role in the decontamination of microbial load from the milk sample. The plasma bubbling generated system efficiently reduced the number of colonies in milk with one month shelf life on 200volt, 10 L/ h, 15 min, 100mL without any negative effect. A non-detrimental effect was observed for physicochemical as well as for nutrient content of plasma bubbling treated milk. To better understand the quality of plasma bubbling on milk, further sensory analysis should be done.

## 6. ACKNOWLEDGEMENTS

The author acknowledges IIFPT, for the technical support and proving infrastructure to carry out the research work. This research did not receive any specific grant from funding agencies in the public, commercial or not-for-profit sectors.

## 7. AUTHORS CONTRIBUTION

Mrs Samarpita Dash conceived the practical work and wrote the manuscript. Dr R. Jaganmohan conceived the manuscript idea. All the authors discussed the methodology and results.

## REFERENCE

1. Ebringer L, Ferenčík M, Krajčovič J. Beneficial health effects of milk and fermented dairy products. Folia microbiol., 2008, 53, 378–394. doi: 10.1007/s12223-008-0059-1.

2. Rowlands A. Bacteriological Standards for Perishable Foods: (c) Milk and Dairy Products. J R Sanit Inst, 1952, 72, 404–410.doi.org/10.1177%2F146642405207200422.

3. National Research Council. Designing foods: animal product options in the marketplace. Natl. Acad. Sci Washington, DC, 1988.

4. Ghodeker DR, Dudani AT, Ranganathan B. Microbiological quality of Indian milk products. J. Milk Food Technol., 1974, 37, 119–122. doi: 10.4315/0022-2747-37.3.119.

5. Camacho AT, Guitian FJ, Pallas E, Gestal JJ, Olmeda S, Goethert H, Spielman A. Serum protein response and renal failure in canine Babesia annae infection. Vet. res, 2005, 36, 713-722. 1. doi: 10.1051/vetres.

6. Lambertini E, Karns J S, Van Kessel J A S, Cao H, Schukken Y H, Wolfgang D R, Pradhan, A K. Dynamics of Escherichia coli virulence factors in dairy herds and farm environments in a longitudinal study in the United States. Appl. Environ. Microobiol, 2015, 81, 4477–4488. doi: 10.1128/AEM.00465-15.

7. Dhanashekar R, Akkinepalli S, Nellutla A. (2012). Milk-borne infections. An analysis of their potential effect on the milk industry. Germs, 2012, 2, 101. doi: 10.11599/germs.2012.1020.

8. Macdonald L E, Brett J, Kelton D, Majowicz S E, Snedeker K, Sargeant J M. A systematic review and meta-analysis of the effects of pasteurization on milk vitamins, and evidence for raw milk consumption and other health-related outcomes. J. Food Prot, 2011, 74, 1814–1832. doi.org/10.4315/0362-028X.JFP-10-269.

9. Datta N, Deeth H C. Diagnosing the cause of proteolysis in UHT milk. LWT-Food Sci. Technol., 2003, 36, 173–182. doi.org/10.1016/S0023-6438(02)00214-1

10. Nair P K, Dalgleish D G, Corredig M. Colloidal properties of concentrated heated milk. Soft Matter, 2013,9, 3815–3824. doi.org/10.1039/C2SM27540F.

11. Ajmal M, Nadeem M, Imran M, Abid M, Batool M, Khan I T, Tayyab M. Impact of immediate and delayed chilling of raw milk on chemical changes in lipid fraction of pasteurized milk. Lipids Health Dis., 2018,17, 1–10. doi: 10.1186/s12944-018-0843-0.

12. Ikawa S, Kitano K, Hamaguchi S. Effects of pH on bacterial inactivation in aqueous solutions due to lowCtemperature atmospheric pressure plasma application. Plasma Processes Polym, 2010, 7, 33–42. doi: 10.1002/ppap.200900090.

13. Kim H J, Yong H I, Park S, Choe W, Jo C. Effects of dielectric barrier discharge plasma on pathogen inactivation and the physicochemical and sensory characteristics of pork loin. Curr Appl Phys, 2013, 13, 1420–1425. doi: 10.1016/j.cap.2013.04.021.

14. Basaran P, Basaran-Akgul N, Oksuz L. Elimination of Aspergillus parasiticus from nut surface with low pressure cold plasma (LPCP) treatment. Food Microbiol., 2008, 25, 626–632. doi: 10.1016/j.fm.2007.12.005.

15. Yu H, Perni S, Shi J J, Wang D Z, Kong M G, Shama G. Effects of cell surface loading and phase of growth in cold atmospheric gas plasma inactivation of Escherichia coli K12. J. Appl. Microbiol, 2006, 101, 1323–1330. doi.org/10.1111/j.1365-2672.2006.03033.x.

16. Kim B, Yun H, Jung S, Jung Y, Jung H, Choe W, Jo C. Effect of atmospheric pressure plasma on inactivation of pathogens inoculated onto bacon using two different gas compositions. Food microbiol, 2011, 28, 9–13. doi: 10.1016/j.fm.2010.07.022.

17. Gurol C, Ekinci F Y, Aslan N, Korachi M. Low temperature plasma for decontamination of E. coli in milk. Int. J. Food Microbiol, 2012, 157, 1–5. doi: 10.1016/j.ijfoodmicro.2012.02.016.

18. Wu X, Luo Y, Zhao F, Mu G. Influence of dielectric barrier discharge cold plasma on physicochemical property of milk for sterilization. Plasma Processes and Polym, 2021, 18, 1900219. doi.org/10.1002/ppap.201900219.

19. Manoharan D, Stephen J, Radhakrishnan M. Study on lowCpressure plasma system for continuous decontamination of milk and its quality evaluation. J. Food Process. Preserv., 2021, 45,0–1. doi: 10.1111/jfpp.15138.

20. Samarpita dash and R. Jaganmohan. ‘Impact Of Plasma Bubbling On Cow Milk: Microbial Reduction And quality Improvement’, Int. J. Life Sci. Pharma Res, 2022, 12, 283–288. doi: 10.22376/ijpbs/lpr.2022.12.1.L283-288.

21. Misra N N, Jo C. Applications of cold plasma technology for microbiological safety in meat industry. Trends Food Sci Technol,2017, 64, 74–86.

22. Surowsky B, Schlüter O, Knorr D. Interactions of non-thermal atmospheric pressure plasma with solid and liquid food systems: a review. Food Eng. Rev., 2015, 7, 82–108.

23. Misra N N, Tiwari B K, Raghavarao K S M S, Cullen P J. Nonthermal plasma inactivation of food-borne pathogens. Food Eng. Rev, 2011, 3, 159–170. doi.org/10.1007/s12393-011-9041-9.

24. Mir S A, Shah M A, Mir M M. Understanding the role of plasma technology in food industry. Food Bioproc Tech, 2016, 9, 734–750. doi.org/10.1007/s11947-016-1699-9.

25. Aparajhitha S, Mahendran R. Effect of plasma bubbling on free radical production and its subsequent effect on the microbial and physicochemical properties of Coconut Neera. Innov Food Sci Emerg Technol.,2019, 58, 102230. doi: 10.1016/j.ifset.2019.102230.

26. Metwally A M, Dabiza N M, El-Kholy W I, Sadek Z I. The effect of boiling on milk microbial contents and quality. J Am Sci, 2011, 7, 110–114.

27. Ray B, Speck M L. Plating procedure for the enumeration of coliforms from dairy products. Appl. Environ Microbiol, 1978, 35, 820–822. doi.org/10.1128/aem.35.4.820-822.1978.

28. Godič Torkar K, Golc-Teger S. ‘The presence of some pathogen microorganisms, yeasts and moulds in cheese samples produced at small dairy-processing plants’, Acta Agric. Slov., 2006, 1, 37–51.

29. Wells J G, Shipman L D, Greene K D, Sowers E G, Green J H, Cameron D N, Ostroff S M. Isolation of Escherichia coli serotype O157: H7 and other Shiga-like-toxin-producing E. coli from dairy cattle. J. Clin. Microbiol, 1991, 29, 985–989. doi.org/10.1128/jcm.29.5.985-989.1991.

30. Marshall V, Poulson-Cook S, Moldenhauer J. Comparative mold and yeast recovery analysis (the effect of differing incubation temperature ranges and growth media). PDA J Pharma Sci Technol, 1998, 52, 165–169.

31. Alves M N, Nesterenko P N, Paull B, Haddad P R, Macka M. Separation of superparamagnetic magnetite nanoparticles by capillary zone electrophoresis using nonCcomplexing and complexing electrolyte anions and tetramethylammonium as dispersing additive. Electrophoresis, 2018, 39, 1429–1436. doi: 10.1002/elps.201800095

32. Muhammad A I, Li Y, Liao X, Liu D, Ye X, Chen S, Ding T. Effect of dielectric barrier discharge plasma on background microflora and physicochemical properties of tiger nut milk. Food Control, 2019, 96, 119–127.doi.org/10.1016/j.foodcont.2018.09.010.

33. Binti Zakaria Z, Yun W S, Alias N, Noor S N M, Mustapha Z, Hussin N, Yusoff N A M. Physicochemical composition, microbiological quality and consumers’ acceptability of raw and pasteurized locally produced goat milk. Mal J. of Fund. Appl. Sci, 2020, 16, 475–482.

34. Cavalcanti A L, Fernandes L V, Barbosa A S,Vieira F F. pH, Titratable Acidity and Total Soluble Solid Content of Pediatric Antitussive Medicines. Acta Stomatol. Croat, 2008, 42.

35. Barbano D M, Lynch J M, Fleming J R. Direct and indirect determination of true protein content of milk by Kjeldahl analysis: collaborative study. J. Assoc. Off. Anal Chem, 1991,74, 281–288. doi: 10.1093/jaoac/74.2.281.

36. Kleyn D H, Lynch J M, Barbano D M, Bloom M J, Mitchell M W, Collaborators: Cooper LS Cusak E Fick M Hanks T Hesen MK Johnson J Kleyn DH Mercer F Monahan D Peat B Petit M. Determination of fat in raw and processed milks by the Gerber method: collaborative study. J. AOAC Int., 2001,84, 1499–1508. doi.org/10.1093/jaoac/84.5.1499.

37. Ting K, Liu Y F, Tian-Li G, Lu-Hua Z. Relationships between viscosity and the contents of macromolecular substances from milk with different storage styles. Food Sci Technol,2016, 4, 49–56. 49–56. doi: 10.13189/fst.2016.040401.

38. Girling P J. Packaging of food in glass containers. Food packaging technology. In: Richard Coles, Dereck Mc Do-well, Mark J Kirwan editors. Food packaging Technology. London, UK: Blackwell Publishing, CRC Press. 2003, 152–73.

39. Sepulveda D R, Góngora-Nieto M M, Guerrero J A, Barbosa-Cánovas G V. Shelf life of whole milk processed by pulsed electric fields in combination with PEF-generated heat. LWT-Food Sci Technol, 2009, 42, 735–739. doi.org/10.1016/j.lwt.2008.10.005

40. Dobrynin D, Fridman G, Friedman G, Fridman A. Physical and biological mechanisms of direct plasma interaction with living tissue. New J. Physiol., 2009, 11, 115020. doi: 10.1088/1367-2630/11/11/115020.

41. Coutinho N M, Silveira M R, Rocha R S, Moraes J, Ferreira M V S, Pimentel T C, Cruz A G. Cold plasma processing of milk and dairy products. Trends in Food Sci. Technol, 2018, 74, 56–68. doi: 10.1016/j.tifs.2018.02.008.

42. Moreau M, Orange N, Feuilloley M G J. Non-thermal plasma technologies: new tools for bio-decontamination. Biotechnol. Adv., 2008, 26, 610–617. doi.org/10.1016/j.biotechadv.2008.08.001.

43. EAS (East African Standards) Raw Cow Milk Specifications. EAS 67:2006, ICS 67.100 HS 04014.20.00 (EAS 67:2000).

44. Foster EM, Nelson FE, Speck ML, Doetsch RN, Olson JC. Dairy microbiology. Prentice-Hall, Inc., Dairy Sci.,1957, 29, 439452. New York.

45. Randolph H E, Chakraborty B K, Hampton O, Bogart D L. Microbial counts of individual producer and commingled grade A raw milk. J. Milk Food Technol, 1973, 36, 146–151. doi.org/10.4315/0022-2747-36.3.146.

46. Jones F T, Langlois B E. Microflora of retail fluid milk products. J. Food prot., 1977, 40, 693–697. doi: 10.4315/0362-028x-40.10.693.

47. Fleet G H, Mian M A. The occurrence and growth of yeasts in dairy products. Int. J. Food Microbiol,1987, 4(2), 145–155. doi: 10.1016/0168-1605(87)90021-3.

48. Fröhlich-Wyder M T. ‘Yeasts in dairy products’, Yeasts in Food, (1970) (2003), 209– 237. doi: 10.1016/B978-1-85573-706-8.50013-7.

49. Tampieri F, Ginebra M P, Canal C. Quantification of plasma-produced hydroxyl radicals in solution and their dependence on the Ph. Analytical chemistry,2021, 93, 3666–3670. doi.org/10.1021/acs.analchem.0c04906.

50. Vasbinder A J, De Kruif C G. Casein–whey protein interactions in heated milk: the influence of pH. Int. Dairy J., 2003, 13, 669–677.doi.org/10.1016/S0958-6946(03)00120-1.

51. Guo J, Huang K, Wang J. Bactericidal effect of various non-thermal plasma agents and the influence of experimental conditions in microbial inactivation: A review. Food Control, 2015, 50, 482–490. doi: 10.1016/j.foodcont.2014.09.037.

52. Raynal-Ljutovac K, Park Y W, Gaucheron F, Bouhallab S. Heat stability and enzymatic modifications of goat and sheep milk. Small Rumin. Res.,2007, 68, 207–220. doi: 10.1016/j.smallrumres.2006.09.006.

53. Prasantha B D, Wimalasiri K M S. Effect of HTST thermal treatments on end-use quality characteristics of goat milk. Int. J. food sci, 2019. doi.org/10.1155/2019/1801724.

54. Ben’Ko E M, Manisova O R, Lunin V V. Effect of ozonation on the reactivity of lignocellulose substrates in enzymatic hydrolyses to sugars. Russ J. Phys. Chem. A, 2013,87, 1108–1113. doi: 10.1134/S0036024413070091.

55. Almeida F D L, Gomes W F, Cavalcante R S, Tiwari B K, Cullen P J, Frias J M, Rodrigues S. Fructooligosaccharides integrity after atmospheric cold plasma and high-pressure processing of a functional orange juice. Food Res. Int, 2017, 102, 282–290. doi: 10.1016/j.foodres.2017.09.072.

56. McAuley, C M, Singh T K, Haro-Maza J F, Williams R, Buckow, R. Microbiological and physicochemical stability of raw, pasteurised or pulsed electric field-treated milk. Innov Food Sci Emerg Technol, 2016, 38, 365–373. doi: 10.1016/j.ifset.2016.09.030.

57. Gavahian M, Chu Y H, Khaneghah A M, Barba F J, Misra N N. A critical analysis of the cold plasma induced lipid oxidation in foods. Trends Food Sci. Technol.,2018, 77, 32–41. doi: 10.1016/j.tifs.2018.04.009.

58. Kim H J, Yong H I, Park S, Kim K, Choe W, Jo C. Microbial safety and quality attributes of milk following treatment with atmospheric pressure encapsulated dielectric barrier discharge plasma. Food Control, 2015, 47, 451–456. doi: 10.1016/j.foodcont.2014.07.053.

59. Elhasan S M, Bushara A M, Abdelhakam K E, Elfaki H A, Eibaid A I, Farahat F H, Sukrab A M. Effect of heat treatments on physico-chemical properties of milk samples. J. Acad. Ind. Res., (JAIR),2017, 6, 40–46.

60. Rolls B A, Porter J W G. Some effects of processing and storage on the nutritive value of milk and milk products. Pro. Nutr Soc,1973, 32, 9–15.

61. Yan D, Cui H, Zhu W, Talbot A, Zhang L G, Sherman J H, Keidar M. The strong cell-based hydrogen peroxide generation triggered by cold atmospheric plasma. Sci. Rep., 2017, 7, 1–9. doi.org/10.1038/s41598-017-11480-x.

62. Cass O W, Paris J P, Stock A M. Research on The Stability of High Strength H2O2. DU PONT DE NEMOURS (EI) AND CO WILMINGTON DE.1966.

63. Liu F, Sun P, Bai N, Tian Y, Zhou H, Wei S, Fang J. Inactivation of bacteria in an aqueous environment by a directCcurrent, coldCatmosphericCpressure air plasma microjet. Plasma Processes Polym, 2010, 7, 231–236. doi: 10.1002/ppap.200900070.

64. Kornacki J L. Enterobacteriaceae, coliforms and Escherichia coli as quality and safety indicators. Compendium of methods for the microbilogical examination of foods, 2001, 69–82.

65. Căpriţă A, Căpriţă R, Creţescu I. The effects of storage conditions on some physicochemical properties of raw and pasteurized milk. J. Agroaliment. Processes. Technol., 2014. 20,198–202.

66. Mittal S, Bajwa U. Effect of heat treatment on the storage stability of low-calorie milk drinks. J. Food Sci. Technol, 2014, 51, 1875–1883. doi.org/10.1007/s13197-012-0714-z.

67. Sarangapani C, Keogh D R, Dunne J, Bourke P, Cullen, P. J. (2017). Characterisation of cold plasma treated beef and dairy lipids using spectroscopic and chromatographic methods. Food Chemistry, 2017, 235, 324–333. doi: 10.1016/j.foodchem.2017.05.016.

68. Wu X, Luo Y, Zhao F, Mu G. Influence of dielectric barrier discharge cold plasma on physicochemical property of milk for sterilization. Plasma Processes and Polym,2021: 18, 1900219. doi: 10.1002/ppap.201900219.

